# *De novo* synthesized polyunsaturated fatty acids operate as both host immunomodulators and nutrients for *Mycobacterium tuberculosis*

**DOI:** 10.1101/2021.07.08.451715

**Authors:** Thomas Laval, Laura Pedró-Cos, Wladimir Malaga, Laure Guenin-Macé, Alexandre Pawlik, Véronique Mayau, Hanane Yahia-Cherbal, Wafa Frigui, Justine Bertrand-Michel, Christophe Guilhot, Caroline Demangel

**Affiliations:** Immunobiology of Infection Unit, Institut Pasteur, INSERM U1221, Paris, France; Université de Paris, Paris, France; Institut de Pharmacologie et de Biologie Structurale (IPBS), Université de Toulouse, CNRS-UPS UMR 5089, Toulouse, France; Integrated Mycobacterial Pathogenomics Unit, Institut Pasteur, CNRS UMR 3525, Paris, France; Immunoregulation Unit, Institut Pasteur, INSERM U122; MetaboHUB-MetaToul, National Infrastructure of Metabolomics and Fluxomics, Toulouse, 31077, France; I2MC, Université de Toulouse, INSERM, Université Toulouse III - Paul Sabatier (UPS), Toulouse, France

## Abstract

Successful control of *Mycobacterium tuberculosis* (Mtb) infection by macrophages relies on immunometabolic reprogramming, where the role of fatty acids (FAs) remains poorly understood. Recent studies unraveled Mtb’s capacity to acquire saturated and monounsaturated FAs *via* the Mce1 importer. However, upon activation macrophages produce polyunsaturated FAs (PUFAs), mammal-specific FAs mediating the generation of key immunomodulatory eicosanoids. Here, we asked whether *de novo* synthesis of PUFAs is modulated in Mtb-infected macrophages and benefits host or pathogen. Quantitative lipidomics revealed that Mtb infection activates the early PUFA biosynthetic pathway for production of eicosanoids. While PUFA synthesis blockade significantly impaired the inflammatory and antimicrobial responses of infected macrophages, it had no effect on Mtb growth *in vivo*. Using a click-chemistry approach, we found that Mtb efficiently imports PUFAs of the ω6 subset via Mce1 in axenic culture, including the eicosanoid precursor arachidonic acid (AA). Notably, Mtb preferentially internalized AA over all other FAs within infected macrophages, but here Mtb’s import of AA was largely Mce1-independent and correlated with elevated AA uptake by host cells. Together, these findings reveal AA as a major FA substrate for intracellular Mtb. They suggest that Mtb’s hijacking of host-derived AA may counteract its stimulatory effect on anti-mycobacterial immune responses.

## INTRODUCTION

*Mycobacterium tuberculosis* (Mtb), the causative agent of human Tuberculosis (TB), caused 1.6 million deaths in 2017 and it is estimated that 23% of the world’s population has a latent TB infection. This success is due to Mtb evolving sophisticated strategies to survive intracellularly in macrophages, its preferred habitat, for long periods of time (Bussi and Gutierrez, 2019). In particular, Mtb’s capacity to import and metabolize host-derived lipids, including fatty acids (FAs) and cholesterol, contributes to long-term persistence *in vivo* (Nazarova et al., 2017, 2019; Pandey and Sassetti, 2008). Recent work indicated that Mtb can acquire saturated and monounsaturated FAs (SFAs and MUFAs, respectively) via a dedicated protein machinery named Mce1, which is coordinated with Mce4-mediated import of cholesterol and plays an important role in Mtb lipid homeostasis (Laval et al., 2021; Lee et al., 2013; Nazarova et al., 2017, 2019; Wilburn et al., 2018).

In addition to SFAs and MUFAs, mammalian cells can produce the additional subset of polyunsaturated FAs (PUFAs), whose secondary metabolites play key immunomodulatory functions in activated macrophages (Dennis and Norris, 2015; Mayer-Barber and Sher, 2015). Interestingly, Toll-like receptors (TLRs) induce signal-specific reprogramming of FA synthesis in macrophages, with differential impact on antibacterial immunity (Hsieh et al., 2020). How Mtb infection modulates PUFA biosynthesis by host macrophages, and whether this enhances inflammation and promotes anti-mycobacterial immune responses is unknown.

Here, we combined quantitative lipidomics with genetic ablation and pharmacological inhibition approaches to assess the importance of the PUFA biosynthetic pathway in innate control of Mtb infection by macrophages. We found that PUFA biosynthesis contributes to the generation of inflammatory and antimicrobial responses of infected macrophages, but this stimulatory effect on macrophage effector functions was not matched by an enhanced capacity to restrict intracellular Mtb infection. This led us to propose that newly generated PUFAs may, like SFAs and MUFAs, serve as FA sources for intracellular Mtb. Our *in vitro* and cellular assays using traceable alkyne-FAs support this hypothesis by showing that Mtb efficiently internalizes the ω6 subset of PUFAs in axenic cultures, and preferentially the eicosanoid precursor arachidonic acid (AA) within macrophages. They also indicated that within macrophages, Mtb’s uptake of FAs is only partially mediated by Mce1, suggesting the existence of alternative import mechanisms *in cellulo*. Together, our findings reveal the pro-inflammatory function of the PUFA biosynthetic pathway during Mtb infection and identify PUFAs as major FA sources for intracellular Mtb.

## RESULTS

### *M. tuberculosis* infection stimulates the biosynthesis of SFAs, MUFAs and upstream PUFAs by host macrophages

Mtb’s impact on host FA metabolism was investigated by infecting bone marrow-derived macrophages (BMDMs), extracting total and free cellular FAs and quantifying each FA species by gas chromatography, with normalization to total DNA. Mtb triggered a significant increase in intracellular levels of free SFA palmitic acid (PA) and MUFAs (oleic and vaccenic acids; OA and VA) after 24h (Fig. 1A-B). These changes were not specific of the Mtb pathogen, as BMDM infection with the *M. bovis* BCG vaccine induced comparable SFA/MUFA profiles (Fig. 1A-B). In both cases, infection-driven variations in free FA levels were associated with parallel trends in total FA levels (Fig. S1A). *De novo* synthesis of SFAs is controlled by the FASN rate-limiting enzyme, and MUFAs are generated from SFAs by the SCD2 rate-limiting enzyme (Fig. 1A) (Guillou et al., 2010). In Mtb-infected BMDMs, increased levels of SFAs were associated with upregulation of *Fasn* transcript levels at 6h post-infection (Fig. 1C). Although *Scd2* mRNA expression remained unchanged (data not shown), the OA:SA ratio reflecting the efficacy of SFA conversion into MUFAs was increased upon Mtb and BCG infection (Fig. 1D).

**Figure 1.**
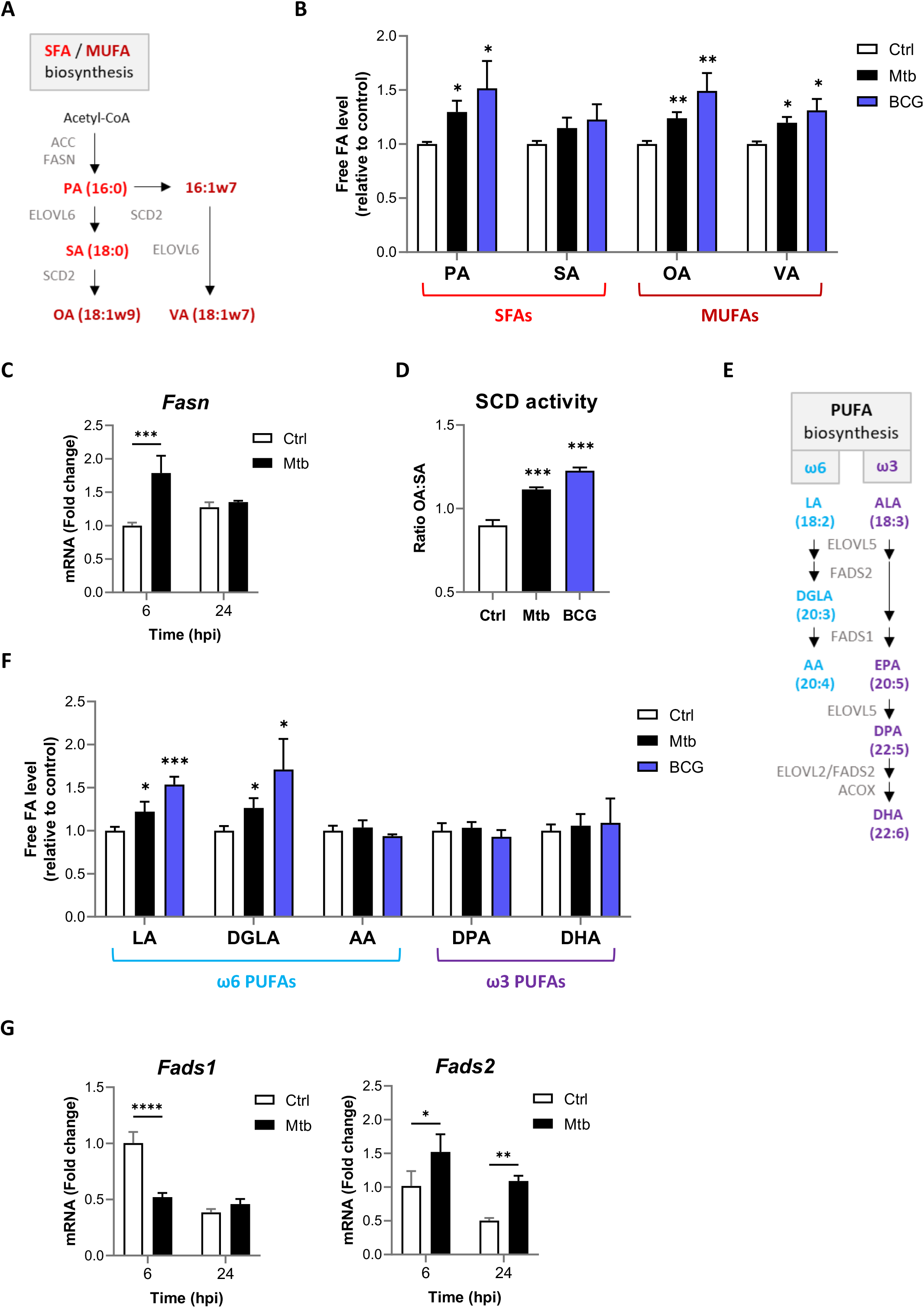
*M. tuberculosis* infection stimulates the biosynthesis of SFAs, MUFAs and upstream PUFAs by host macrophages. (**A**) Schematics of SFA and MUFA biosynthetic pathways. PA: palmitic acid, SA: stearic acid, OA: oleic acid, VA: vaccenic acid. (**B**) Intracellular levels of free SFAs and MUFAs in BMDMs infected with *M. bovis* BCG (BCG) or *M. tuberculosis* H37Rv (Mtb) at the same multiplicity of infection (MOI) of 2:1 for 24h. FA levels were normalized to total DNA content, and are shown as fold change relative to Ctrl. (**C**) Relative mRNA expression of *Fasn* in BMDMs infected with Mtb for the indicated times, as determined by qRT-PCR. **(D)** SCD activity in BMDMs treated as in (**B**), as estimated by the ratio of OA to SA levels. (**E**) Schematics of PUFA biosynthetic pathways. LA: linoleic acid, DGLA: dihomo-γ-LA, AA: arachidonic acid, ALA: α-linolenic acid, DPA: docosapentaenoic acid, DHA: docosahexaenoic acid. (**F**) Intracellular levels of free PUFAs in BMDMs treated as in (**B**). (**G**) Relative mRNA expression of *Fads1* and *Fads2* in BMDMs infected with Mtb for the indicated times, as determined by qRT-PCR. All data are means +/- SD (n=3) and are representative of 2 independent experiments. \**P*<0.05, \*\**P*<0.01, \*\*\**P*<0.001, \*\*\*\**P*<0.0001, unpaired Student’s t-tests [(B), (D) and (F)] and two-way ANOVA with Bonferroni post-hoc multiple comparison tests [(C) and (G)].

With regard to PUFAs, we detected an increased level of ω6 precursor linoleic acid (LA) that was associated with elevated levels of its conversion product dihomo-gamma-linolenic acid (DGLA), but not the DGLA product AA (Fig. 1E-F). On the ω3 PUFA side, alpha-linolenic acid (ALA) and eicosapentaenoic acid (EPA) were below detection limit, and the long chain ω3 PUFAs docosapentaenoic acid (DPA) and docosahexaenoic acid (DHA) were not modulated by Mtb infection (Fig. 1E-F). Again, BMDM infection with *M. bovis* BCG induced comparable profiles of free and total PUFAs (Fig. 1F and S1B). Conversion of ω6 and ω3 PUFA precursors into downstream PUFAs is jointly controlled by 3 enzymes: the FA desaturases (FADS)1 and FADS2, and the elongase ELOVL5 (Fig. 1E). In BMDMs infected with Mtb, *Fads1* transcript levels were repressed 6h post-infection, while those of *Fads2* and *Elovl5* were increased and unchanged, respectively (Fig. 1G and unshown data). Transcriptional induction of *Fads2* combined to repression of *Fads1* was thus consistent with the observed accumulation of LA and DGLA, but not AA, DPA and DHA PUFAs in Mtb-infected BMDMs. Together, our data shown in Figure 1 demonstrated that Mtb infection upregulates the biosynthesis of SFAs, MUFAs and upstream, but not downstream PUFAs in host macrophages.

### FADS2 inhibition impairs the effector functions of macrophages during *M. tuberculosis* infection

Long chain PUFAs can be mobilized by hydrolysis of phospholipids to fuel the production of lipid mediators of inflammation (Dennis and Norris, 2015). In particular, conversion of AA by the COX/LOX pathways generates eicosanoids (Fig. 2A) modulating the ability of macrophages to control mycobacterial infection (Mayer-Barber and Sher, 2015). Since the early PUFA biosynthetic pathway was activated by Mtb infection through transcriptional induction of *Fads2*, we wondered if enough AA was neosynthesized to mediate the generation of eicosanoids, despite the repression of *Fads1* expression. To test this, we used a selective inhibitor of FADS2 (SC-26196, hereafter named iFADS2) (Obukowicz et al., 1998). Treating BMDMs with iFADS2 induced a dose-dependent defect in macrophage ability to convert deuterated LA into AA (Fig. 2B), with complete inhibition of FADS2 activity observed in the presence of 5 µM inhibitor. Consistent with previous studies (Chen et al., 2008; Knight et al., 2018), BMDMs infected with Mtb secreted higher amounts of AA metabolites deriving from both the COX (Fig. 2C) and LOX pathways (Fig. 2D), compared to non-infected controls. However, when BMDMs were infected with Mtb in the presence of 5 µM iFADS2, the production of AA metabolites generated by COX and LOX was significantly reduced (Fig. 2C-D), demonstrating that infection-induced eicosanoids largely originate from *de novo* synthesized PUFAs.

**Figure 2.**
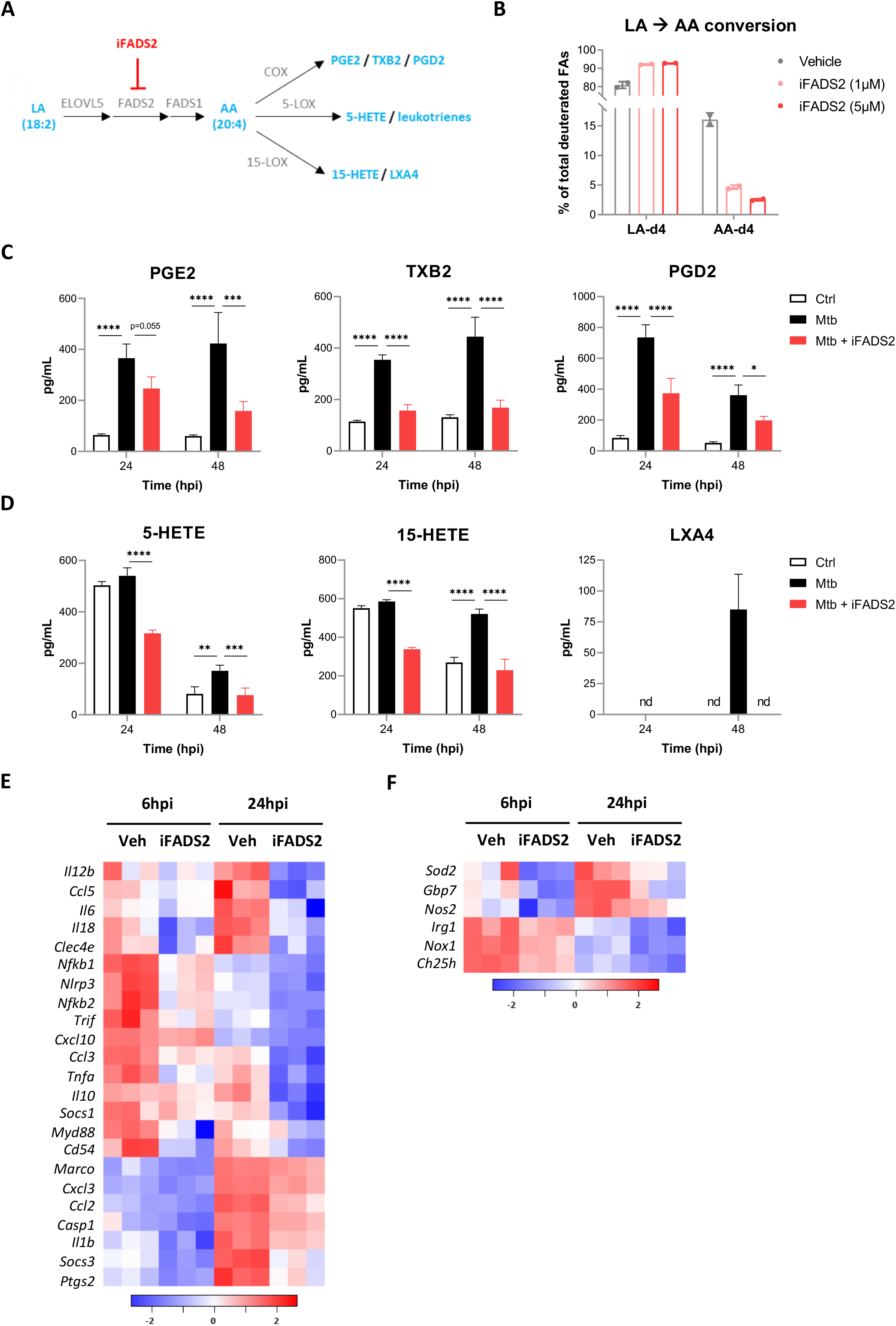
FADS2 inhibition impairs the effector functions of macrophages during *M. tuberculosis* infection. (**A**) Schematics of biosynthetic pathways of AA-derived eicosanoids. COX: cyclooxygenase, LOX: lipoxygenase, PG: prostaglandin, TX: thromboxane, HETE: hydroxyeicosatetraenoic acid, LX: lipoxin. (**B**) Intracellular levels of deuterated ω6 PUFAs in BMDMs treated with indicated doses of FADS2 inhibitor (iFADS2) or vehicle control, and incubated with the ω6 precursor LA-d4 for 24h, relative to total deuterated FAs levels. Data are means +/- SD (n=2). (**C-D**) Secreted levels of COX- (**C**) and LOX-derived (**D**) metabolites of AA by BMDMs either uninfected (Ctrl), or infected with Mtb and treated with 5µM of FADS2 inhibitor (Mtb + iFADS2) or with vehicle control (Mtb) for 24 or 48h. Data are means +/- SD (n=3), \**P*<0.05, \*\**P*<0.01, \*\*\**P*<0.001, \*\*\*\**P*<0.0001 in a two-way ANOVA with Dunnett post-hoc multiple comparison tests. (E-F) Heatmap of mRNA expression levels of inflammatory (E) and antimicrobial (F) genes determined by NanoString analysis of BMDMs infected with Mtb, and treated with 5µM of iFADS2 or vehicle control (Veh) for 6 or 24h. Shown are genes that were significantly downregulated by iFADS2 treatment at 6 and/or 24h (fold change of at least 1.15 and FDR<0.05, two-way ANOVA with Benjamini-Hochberg adjustment for multiple comparison). Source data are available in Figure 2–source Data 1: Complete list of normalized mRNA levels in BMDMs non-infected (NI Ctrl), or infected with Mtb and treated with iFADS2 (Mtb iFADS2) or vehicle control (Mtb Veh), as determined by NanoString analysis.

To determine if FADS2 inhibition alters the innate immune functions of macrophages, we profiled the expression of a panel of genes involved in antimicrobial and inflammatory responses using a custom NanoString nCounter CodeSet (Table S1). BMDMs were infected with Mtb and treated with 5 µM iFADS2, and gene expression was assessed at 6 and 24h post-infection. FADS2 inhibition induced a significant downregulation of major inflammatory genes (*Tnfa*, *Il1b*, *Il6*) (Fig. 2E) and genes involved in antimicrobial responses of macrophages (*Nos2*, *Nox1*, *Irg1*, *Sod2*) (Fig. 2F). Overall, our lipidomic and transcriptional analyses of FADS2-inhibited macrophages indicated that, despite the transcriptional repression of *Fads1*, Mtb-driven activation of the PUFA biosynthetic pathway fuels the production of eicosanoids and promotes the antimicrobial and inflammatory responses of host macrophages.

### Blocking host PUFA biosynthesis does not impact *M. tuberculosis* growth *in vivo*

Our data in Figure 2 predicted that FADS2 inhibition should limit macrophage capacity to restrict the intracellular growth of Mtb. To test this hypothesis, we infected BMDMs with Mtb in the presence or absence of iFADS2, and monitored mycobacterial growth during 5 days by titrating colony-forming units (CFUs) in cell lysates. Neither pharmacological inhibition of FADS2 nor siRNA knock-down of FADS2 (Fig. S2A) had a detectable effect on intracellular growth of Mtb in BMDMs (Fig. 3A-B). To determine if a complete defect in FADS2 activity was necessary to impact Mtb intracellular growth, we knocked out FADS2 in the THP-1 human cell line using the CRISPR-Cas9 approach. We selected 3 independent clones (bearing distinct point mutations in exon 2 of *FADS2* gene) that lacked FADS2 protein expression (Fig. S2B) and showed a severe defect in PUFA conversion (Fig. S2C). In line with our data using iFADS2 and siRNAs, Mtb grew similarly in all tested wild-type (WT) and knock-out (KO) clones (Fig. 3C).

**Figure 3.**
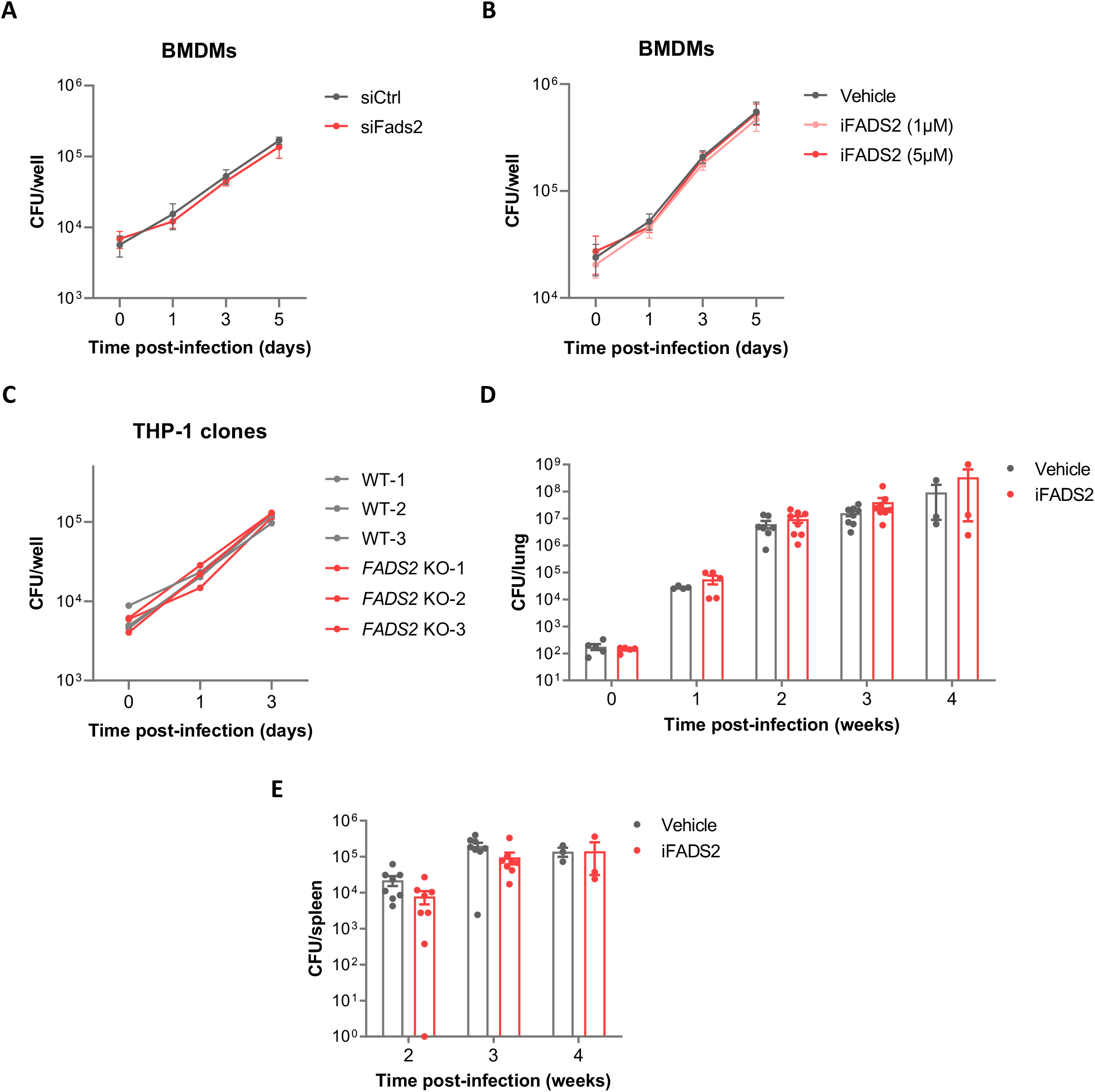
Blocking host PUFA biosynthesis does not impact *M. tuberculosis* growth *in vivo*. (**A-B**) Intracellular growth of Mtb inside BMDMs transfected with non-targeting (siCtrl) or *Fads2* siRNA (siFads2) (**A**), or treated with a FADS2 inhibitor (iFADS2), or with vehicle control (**B**), as determined by CFU plating at the indicated days post-infection. All BMDM infection data are means +/- SD (n=3) and are representative of at least 2 independent experiments. (**C**) Intracellular growth of Mtb inside THP-1 wild-type (WT) or *FADS2* knockout (KO) clones, as determined by CFU plating at the indicated days post-infection. Each line represents mean CFUs (n=3) in one independent THP-1 clone. (**D-E**) Growth of Mtb in the lungs (**D**) and spleen (**E**) of mice treated with iFADS2 or with vehicle control during 4 weeks, as determined by CFU plating. Shown data are means +/- SEM of 2 pooled independent experiments (Exp1: n=3, Exp2: n=5 mice per timepoint).

Since several lipid mediators and inflammatory genes modulated by iFADS2 are involved in adaptive immunity against Mtb (Mayer-Barber and Sher, 2015), we investigated the effect of a systemic inhibition of FADS2 in a mouse model of Mtb aerosol infection. In the conditions tested, mouse treatment with a FADS2 inhibitor did not significantly alter Mtb growth in lungs and spleen (Fig. 3D-E). Therefore, although it impairs innate immune responses of macrophages known to play important roles in Mtb infection outcome, blocking PUFA biosynthesis does not impact Mtb growth *in vivo*.

### *M. tuberculosis* efficiently imports ω6 PUFAs through the Mce1 transporter in axenic culture

We hypothesized that the anti-mycobacterial effects of PUFA biosynthesis on macrophage effector functions could be masked by a pro-mycobacterial role of PUFAs as nutrient sources for Mtb. Indeed, SFAs and MUFAs like PA and OA have been shown to be readily imported and metabolized by Mtb in axenic cultures and within macrophages (Nazarova et al., 2017, 2019). To determine if Mtb has ability to import PUFAs, we used alkyne-FAs, structural analogs of natural FAs that can be detected by click-chemistry using azide-fluorochromes (Thiele et al., 2012) (Fig. 4A). Using this approach, we confirmed that PA, and OA to a lower extent, are efficiently internalized by Mtb (Fig. 4B). While ω3 PUFAs (EPA and DHA) uptake was barely detected, we found that ω6 PUFAs (LA and AA) were imported by Mtb as efficiently as OA.

**Figure 4.**
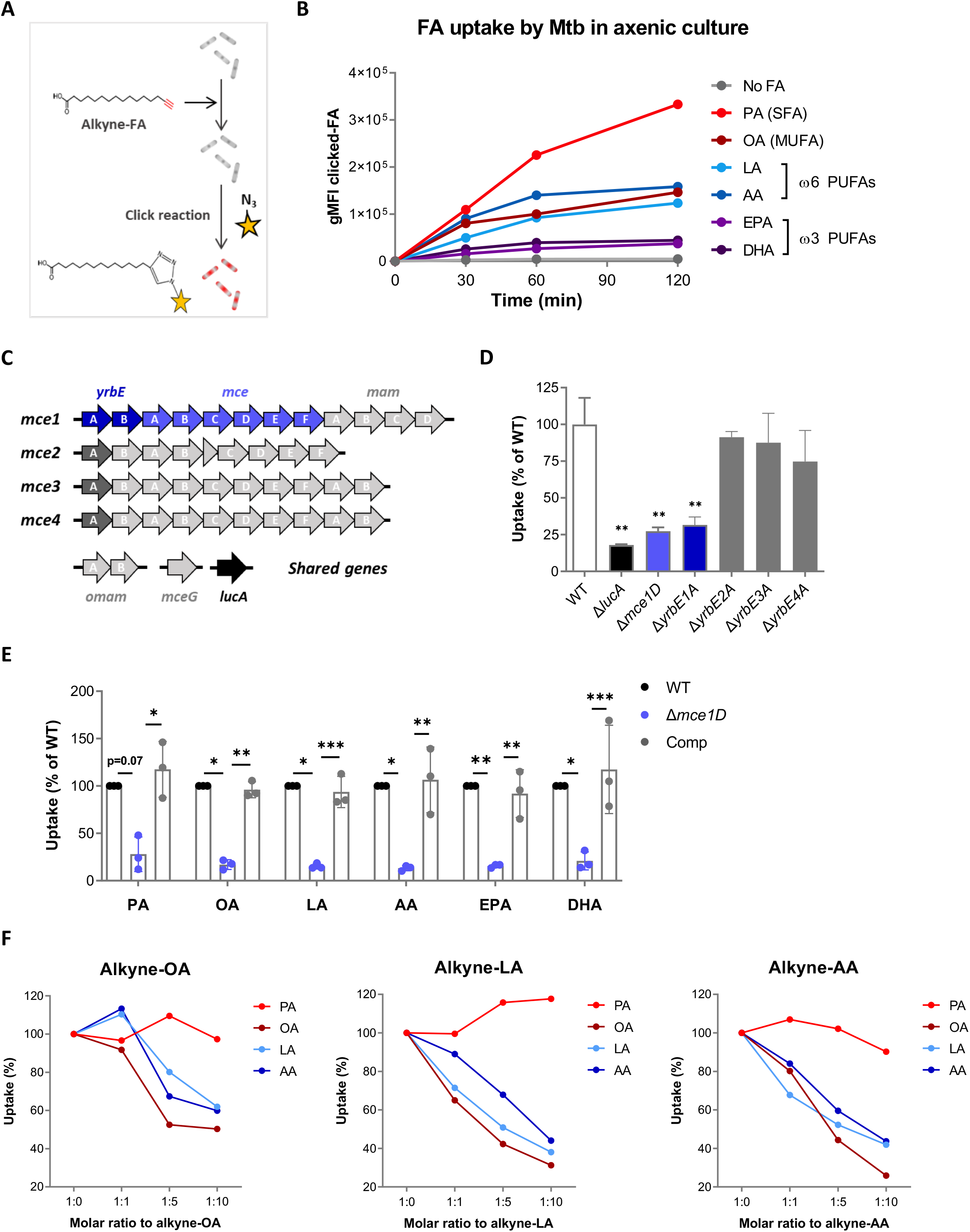
*M. tuberculosis* efficiently imports ω6 PUFAs through the Mce1 transporter in axenic culture. (**A**) Schematics of the click-chemistry approach used to compare the uptake of SFAs, MUFAs and PUFAs by Mtb in axenic culture. (**B**) Differential kinetics of alkyne-SFA, -MUFA and -PUFA uptake (all added at a concentration of 20µM) by Mtb, as estimated by flow cytometry assessment of the geometric mean fluorescence intensities (gMFI) of clicked-FA. Shown data are representative of 3 independent experiments. (**C**) Schematics of the composition of the *mce1-4* operons and related genes in Mtb genome. (**D**) Uptake of alkyne-PA by different Mtb strains deficient for the expression of genes belonging to *mce1-4* operons, relative to wild-type (WT) Mtb. Data are means +/- SD (n=3), representative of 2 independent experiments, \*\**P*<0.01, unpaired Student’s t-tests. (**E**) Uptake of alkyne-FAs by Mtb Δ*mce1D* and its complemented counterpart (Comp), relative to WT. Data are means +/- SD from 3 independent experiments, \**P*<0.05, \*\**P*<0.01, \*\*\**P*<0.001, paired t-tests. (**F**) Relative uptake of alkyne-OA, -LA and -AA by Mtb WT, in the presence of increasing amounts of natural PA, OA, LA or AA. Shown data are representative of at least 2 independent experiments.

We next assessed the role of Mce1 as importer of PUFAs in Mtb, by introducing inactivating mutations in genes of Mce operons by allelic exchange (Fig. 4C and S3A). In accordance with previous studies (Nazarova et al., 2017, 2019), knocking out *lucA, mce1D* or *yrbE1A*, but not *yrbE2A, yrbE3A and yrbE4A* resulted in pronounced defects in alkyne-PA and -OA import (as shown for PA in Fig. 4D). Likewise, we found that Mtb’s import of alkyne-LA, -AA, -EPA and -DHA was abrogated in the *ΔyrbE1A* (Fig. S3B) and *Δmce1D* mutants (Fig. 4E), and restored in the *Δmce1D::comp* complemented strain (Fig. 4E). This established that Mtb has ability to import all PUFAs via the Mce1 transporter.

Since ω6 PUFAs (LA and AA) were efficiently imported by Mtb via Mce1, we asked whether they competed with other FAs for transport. To test this, we measured Mtb’s uptake of alkyne-FAs as above, in the presence of increasing amounts of natural FAs. Uptake of alkyne-OA, -LA and -AA was dose-dependently decreased by addition of their natural counterpart, validating the conditions of the assay (Fig. 4F). Interestingly, LA and AA competed with each other and with OA, but not with PA (Fig. 4F). Together, data in Figure 4 indicated that SFAs, MUFAs and ω6 PUFAs are imported by Mtb via Mce1 in axenic culture, with variable efficacy. They suggested that ω6 PUFAs compete with OA, but not PA, for uptake by Mce1.

### *M. tuberculosis* preferentially internalizes AA in the context of macrophages

We next investigated if all FAs had comparable ability to traffic to Mtb within infected macrophages. Here, BMDMs were infected with a GFP-expressing strain of Mtb prior to a pulse/chase with equivalent concentrations of alkynes derivatives of PA, OA, LA, AA, EPA or DHA (Fig. S4A). Intra-phagosomal Mtb was then isolated from infected cells, and FAs imported by Mtb were stained by click reaction and quantified by flow cytometry (Fig. S4B). All alkyne-FAs could be detected in intracellular Mtb after 24h, and at lower levels after 72h (Fig. 5A). However, differently from *in vitro* grown Mtb (Fig. 4B), intracellular Mtb showed a clear preference for AA over all other FAs including PA (Fig. 5A). This observation was confirmed by confocal microscopy analysis of alkyne-AA signals in Mtb-infected cells (Fig. 5B), and quantification of alkyne-PA and -AA signals in phagocytozed Mtb (Fig. 5C). It is interesting to note that increased uptake of AA by intracellular Mtb correlated with a relatively higher internalization of this FA by host macrophages, which was independent of Mtb infection (Fig. 5D). Similar differences in cellular and bacterial uptake of PA and AA were observed in human THP-1 macrophages (data not shown). We concluded that preferential uptake of AA by intracellular Mtb reflects the higher bioavailability of this FA in host macrophages.

**Figure 5.**
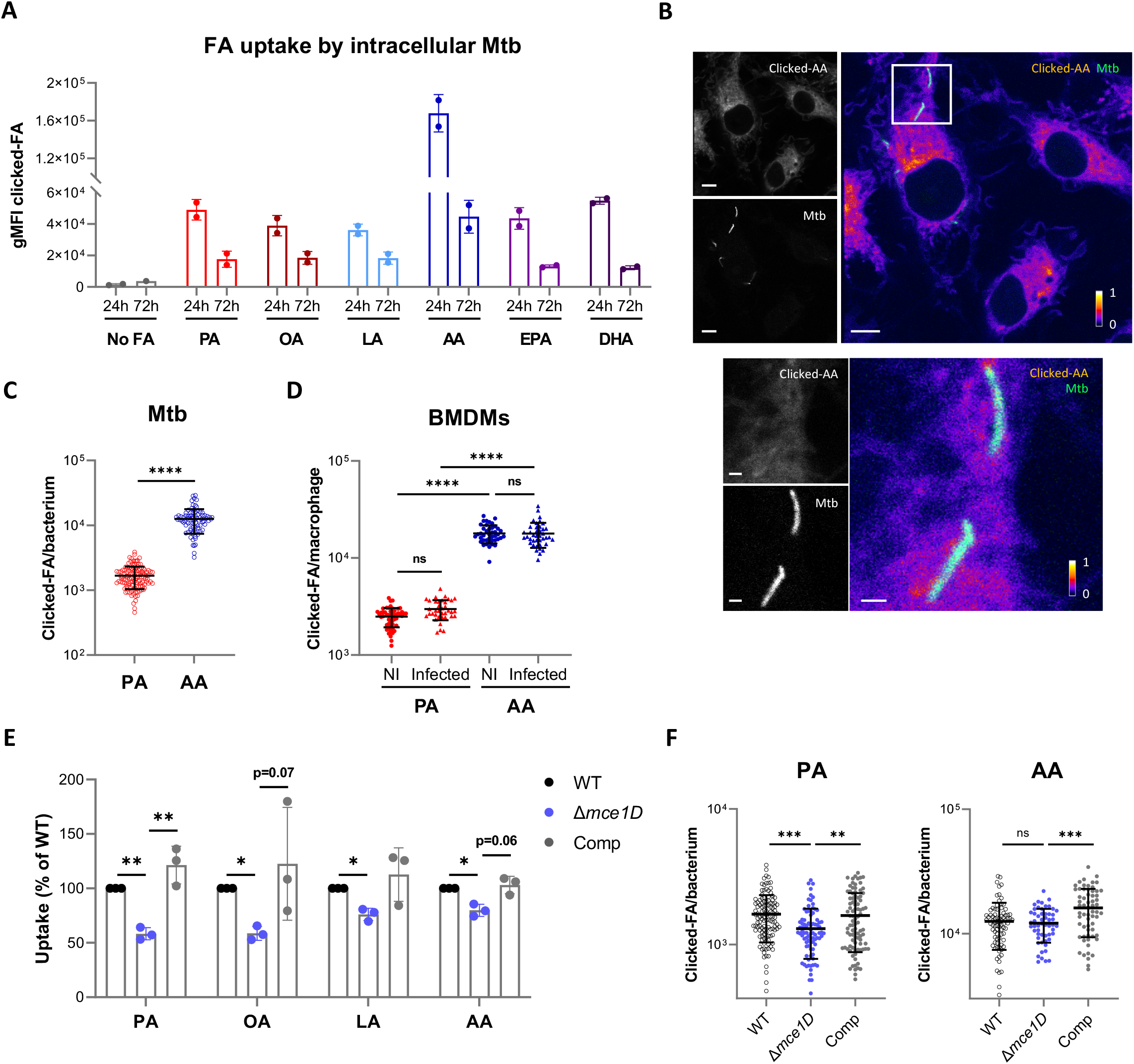
*M. tuberculosis* preferentially internalizes AA in the context of macrophages. (**A**) Differential uptake of alkyne-FAs by Mtb isolated from BMDMs infected for 24 or 72h, as measured by flow cytometry. Data are means +/- SD from 2 independent experiments. (**B**) Distribution of alkyne-AA in Mtb-infected BMDMs at 24h post-infection, as shown on representative confocal images (green = GFP-expressing Mtb, log scale colormap = clicked-AA). Color bar indicates the relative range of pixel intensity (white=high, purple=low, from 0 arbitrary unit to 1). Bar scale=5µm. Enlargement of the boxed area in the merged image (bar scale=1 µm). (**C-D**) Quantification of the alkyne-FA signal in intracellular bacteria detected based on GFP signal (**C**) and in whole BMDMs either non-infected (NI) or Mtb-infected (**D**), using confocal images of BMDMs infected for 24h with a GFP-expressing strain of Mtb. Bars show means +/- SD, n>82 for (C) and n>37 for (D). ns = not significant, \*\*\*\**P*<0.0001 in an unpaired Student’s t-test (C) and one-way ANOVA with Tukey post-hoc multiple comparison tests (D). (E) Uptake of alkyne-FAs by GFP-expressing Mtb Δ*mce1D* and its complemented counterpart (Comp), recovered from BMDMs infected for 24h, relative to GFP-expressing Mtb WT, as analyzed by flow cytometry. Data are means +/- SD from 3 independent experiments, \**P*<0.05, \*\**P*<0.01, paired t-tests. (E) Quantification of the alkyne-FA signal in intracellular bacteria detected based on GFP signal, using confocal images of BMDMs infected for 24h with different GFP-expressing Mtb strains. Bars show means +/- SD, n>54. ns = not significant, \*\**P*<0.01, \*\*\**P*<0.001 in a one-way ANOVA with Dunnett post-hoc multiple comparison tests.

Finally, we assessed the role of Mce1 in PUFA uptake by intracellular Mtb, using GFP-expressing versions of Mtb *Δmce1D* and *Δmce1D::comp*. The uptake of alkyne-FAs by intracellular Mtb was quantified by flow cytometry analysis of isolated bacteria (Fig. 5E), and confocal microscopy analysis of phagocytozed bacteria within infected cells (Fig. 5F). In agreement with previous studies (Nazarova et al., 2019), Mtb’s import of PA and OA was significantly decreased in the absence of Mce1D and restored by gene complementation during macrophage infection (Fig. 5E-F), although to a limited extent compared to Mtb grown in axenic cultures (Fig. 4E). Likewise, Mtb’s import of ω6 PUFA LA and AA was significantly reduced by *mce1D* gene knock-out, but only by 24 and 20%, respectively (Fig. 5E-F). Together, these data indicated that in the context of macrophages, Mtb preferentially imports AA over other host-derived FAs, via mechanisms that are largely independent of the Mce1 transporter. They argue that host PUFAs, and particularly AA, serve as nutrient sources for intracellular Mtb.

## DISCUSSION

In the present study, we show that PUFAs, a host-specific subset of FAs, are imported by Mtb *in vitro* and within macrophages. These findings are consistent with previous observations that radiolabeled LA and ALA were incorporated into cell wall lipids of Mtb grown under axenic conditions (Morbidoni et al., 2006), and that AA used as a sole carbon source was able to support Mtb growth *in vitro* (Forrellad et al., 2014). Our data in Figure 4 indicate that ω6, but not ω3 PUFAs, are imported by Mtb in axenic culture. ω6 PUFAs were imported as efficiently as MUFAs, and competed with MUFAs for Mce1-mediated import in Mtb. This finding suggests that MUFAs and ω6 PUFAs share the same binding site on the Mce1 transporter, or use a common accessory FA-binding protein that directs these FAs to the Mce1 transporter. Based on Mce1 gene inactivation studies (Nazarova et al., 2019; Wilburn et al., 2018), Mce1A-F proteins are believed to form a channel recognizing FA substrates and shuttling them across the mycobacterial cell wall, to deliver them to the putative YrbE1A/B permeases embedded in the cytoplasmic membrane. Our observation that mutations in the *mce1D* and *yrbE1A* genes reduce Mtb’s import of PUFAs by more than 80% *in vitro* supports the view that these proteins play a critical role in recognition and internalization of PUFAs, similar to SFAs and MUFAs. In comparison to all other FAs, ω3 EPA and DHA were poorly internalized by Mtb *in vitro*. However, Mtb’s uptake of ω3 PUFAs also depended on Mce1 in axenic cultures. The distinctive conformational properties of natural, cis ω3 PUFAs, such as high flexibility, may account for a lower affinity for the Mce1 receptor (Chu and Wang, 2014). OA, LA and AA competed with each other for transport by Mce1, but none of these FAs did compete with PA. This suggests that Mce1 displays at least two distinct FA binding sites specialized in recognition of SFAs and MUFAs/PUFAs respectively, a property that may help Mtb optimize the uptake of host-derived FAs during infection.

Our lipidomic analyses of BMDMs infected with Mtb revealed an increase in intracellular levels of free SFAs and MUFAs that was preceded by an upregulation of the gene expression of *Fasn*, the SFA-synthetizing enzyme. We also observed that Mtb represses the gene expression of *Pparg* and several PPARγ targets involved in FA catabolism (*Cpt1a*, *Acox3*, *Lipa*), which may further promote free SFA/MUFA accumulation in infected macrophages (Fig. S5A). Recent studies have identified the MyD88 signaling pathway as a major driver of SFA/MUFA metabolism reprogramming in TLR-stimulated macrophages (Hsieh et al., 2020; Oishi et al., 2017). When we stimulated BMDMs with TLR2/4 agonists, levels of free and total SFAs/MUFAs were both increased (Fig. S6A-B), supporting the hypothesis that Mtb-driven changes in SFA/MUFA synthesis in host macrophages result from recognition of mycobacteria pattern by TLR2/4. Likewise, total LA levels, including free and esterified LA, were upregulated by Mtb infection or TLR2/4 stimulation (Fig. S6C-D). We hypothesize that increased LA levels result from an enhanced uptake of this essential ω6 PUFA precursor by host macrophages upon Mtb infection.

While *de novo* synthesis of SFAs and MUFAs was globally upregulated by Mtb, that of PUFAs was transcriptionally repressed at the level of *Fads1*. Our findings are corroborated by a recent lipidomic study that detected a comparable blockade in long PUFA production by BMDMs stimulated with TLR2 or TLR4 agonists (Hsieh et al., 2020). A transient downregulation of *Fads1* and/or *Fads2* was also reported in a separate study using mouse peritoneal macrophages stimulated with a TLR4 agonist (Oishi et al., 2017). Downregulation of *Fads1* is surprising as *Fasn* was upregulated by Mtb. Both genes are targets of the LXR and SREBP-1 regulators of FA metabolism (Daemen et al., 2013; Joseph et al., 2002). In Mtb-infected BMDMs, upregulation of *Fasn* gene expression was associated with transcriptional induction of *Lxra*, *Ch25h*, and *Srebf1* to a lower extent (Fig. S5B), indicating that Mtb activates LXRs. How *Fads1* transcript levels are concomitantly downregulated remains an open question.

An important new discovery was that Mtb imports AA at significantly higher levels, compared to all other tested FAs, in macrophages pulsed with exogenous FAs. In agreement with previous work by Nazarova *et al*. using macrophages pulsed with Bodipy-PA (Nazarova et al., 2019), *mce1D* contributed to about 40% of total PA uptake by intracellular Mtb (Fig. 5E). In comparison, *mce1D* deficiency had even less of an effect on the uptake of ω6 PUFAs, especially AA. These findings reveal the limited importance of Mce1 in uptake of host FAs by intracellular Mtb, and points to alternative FA import mechanisms. In particular, our observation that the differential uptake of PA and AA by intracellular Mtb correlates with the relative abundance of these FAs in the macrophage cytosol suggests that FA bioavailability in host macrophages is a determining factor. Although intracellular AA levels were unchanged in Mtb-infected BMDMs (Fig. 1F and S1), those of upstream ω6 PUFAs (Fig. 1F and S1) and downstream eicosanoid products (Fig. 2C-D) were upregulated by Mtb infection, suggesting an increased biosynthesis of AA and flux in the COX/LOX pathways. Together with our observation that exogenous AA is superiorly imported by macrophages (Fig. 5B,D) compared to other FAs, our findings support the view that AA represents a major source of FA for intracellular Mtb.

The AA pathway has recently emerged as a potential target of host-directed therapeutic strategies for TB (Young et al., 2020). Catabolism of AA by cyclooxygenase (COX) and lipoxygenase (LOX) yields the eicosanoid products prostaglandins and leukotrienes respectively, which are important signaling molecules modulating inflammation and cell death. While the pro-inflammatory action of eicosanoids is beneficial to the host at early stages of infection, by promoting the intracellular killing of Mtb via induction of tumor necrosis factor α in infected macrophages, it may drive immunopathology at later stages of disease (reviewed in (Young et al., 2020)). Non-steroidal anti-inflammatory drugs inhibiting COX activity are being evaluated in humans as adjunctive therapies to standard TB treatments, with the rationale to limit inflammation-induced tissue pathology. In our study, transcriptional repression of *Fads1* in Mtb-infected macrophages did not prevent the production of AA-derived secondary metabolites, especially the pro-inflammatory COX products PGE2, PGD2 and TXB2 (Fig. 2). In immunopathologic conditions, fully blocking the ω6-PUFA biosynthetic pathway with iFADS2 may thus increase the efficacy of COX inhibitors, while depriving Mtb of additional FA sources.

In conclusion, this study reveals that *de novo* synthesized PUFAs operate both as host immunomodulators and nutrients for Mtb. It suggests that Mtb adapts its FA uptake to the intracellular microenvironment, to take advantage of the AA surge triggered by infection.

## MATERIALS AND METHODS

### Reagents

All solvents for lipid extractions, Phorbol 12-myristate 13-acetate (PMA), fatty acid-free bovine serum albumin (FA-free BSA), ethyleneglycol-bis(β-aminoethyl)-N,N,Nʹ,Nʹ-tetraacetic acid (EGTA), methylcellulose (Methocel A4M), tyloxapol, Tween 20 and Triton X-100 were from Sigma-Aldrich. Ultrapure LPS from *E. coli* serotype 055:B5 TLR grade was purchased from Enzo Life Sciences. Pam3Csk4 was obtained from Invivogen. The FADS2 inhibitor SC-26196 (iFADS2) was purchased from Cayman Chemical, and stored as DMSO stock solutions at -20°C. All natural, deuterated and alkyne derivatives of FAs were obtained from Cayman Chemical, and stored as ethanol stock solutions at - 20°C. Before medium supplementation, FAs were conjugated to albumin by a preincubation with FA-free BSA at a molar ratio of 2:1 (FA:Albumin) 20min at 37°C, unless otherwise stated.

### Macrophage generation and culture conditions

Bone marrow was isolated by perfusion of femurs and tibias from male 7- to 12-week-old C57BL/6J mice (Charles River Laboratories), and cultured in Dulbecco’s modified Eagle’s medium (DMEM, Gibco Laboratories) with 10% heat-inactivated fetal bovine serum (FBS, PAA), 2mM GlutaMAX, 15% L929-conditioned medium, 100U/mL penicillin and 100µg/mL streptomycin (Pen-Strep) for 6 days, with media change on day 3. After differentiation, BMDMs were cultured in DMEM with 10% FBS, 2mM GlutaMAX, and 5% L929-conditioned medium (BMDM media). For *Fads2* knockdown, BMDMs were transfected after 6 days of differentiation with 25nM ON-TARGETplus SMARTpool Fads2 siRNA (Dharmacon) or the same concentration of non-targeting siRNA pool (siCtrl) using lipofectamine RNAiMax (Thermo Fisher Scientific) according to the manufacturer’s instructions. 24h post-transfection, the medium was changed to BMDM media, and cells were used at 48h post-transfection. The THP-1 cell line was obtained from ATCC (American Type Culture Collection, TIB-202) and cultured in RPMI-1640 (Gibco Laboratories) supplemented with 10% FBS, 2mM GlutaMAX (THP-1 media) plus Pen-Strep. THP-1 monocytes were differentiated by incubation with 20ng/mL PMA in THP-1 media for 48h, then PMA was removed and cells were maintained in THP-1 media for an additional 24h before infection.

### CRISPR/Cas9-mediated *FADS2* knockout in THP-1

The guide RNA targeting the exon 2 of *FADS2* gene (5ʹ-GCACCCTGACCTGGAATTCGT-3ʹ) was designed using http://crispor.tefor.net/, and cloned into pSpCas9(BB)-2A-GFP (PX458, Addgene), according to the published protocol (Ran et al., 2013) . THP-1 cells were transfected with the plasmid using the Amaxa Nucleofector 2b device (Lonza) and the Human Monocyte Nucleofector Kit (Lonza) as previously described (Schnoor et al., 2009). The following day, GFP-positive transfected cells were sorted in 96-well plates at a density of 1 cell per well using a FACS ARIA III (BD Biosciences) and cultivated in THP-1 media. Growing cell clones were then screened by PCR using AmpliTaq master mix (Life Technologies) and specific primers (Fwd, 5’-GCACATTTCCAGTGCCAAGG-3’; Rev, 5’-GGAGAGAGGAGACGCCACTA-3’), and by FADS2 immunoblots as described below.

### Mycobacterial growth conditions

The H37Rv strain of Mtb and the Pasteur 1173P2 strain of *M. bovis* BCG and their DsRed-expressing, hygromycin-resistant counterparts were kindly provided by Laleh Majlessi (Insitut Pasteur, Paris) and Nathalie Winter (INRA, Tours), respectively. Bacteria were routinely grown at 37°C in Middlebrook 7H9 broth (BD Bsiosciences) supplemented with 10% BD BBL Middlebrook OADC enrichment (BD Biosciences) and 0.05% tyloxapol. Kanamycin 40µg/mL, hygromycin 50µg/mL, streptomycin 25µg/mL and zeocin 50µg/mL were used for selection.

### Mtb mutant construction and growth conditions

For each target gene, allelic exchange substrates (AES) were constructed by generating PCR fragments with primers X1-X2 and X3-X4, genomic DNA from *M. tuberculosis* H37Rv and enzyme PrimeSTAR GXL (Takara Bio). Such primers allowed introduction of DraIII or Van91I restriction sites (in red and green, respectively in Table S2) at both ends of the PCR fragment. In parallel, the kanamycin (km) resistance cassette was amplified from plasmid pET26b (Novagen) using primers k1 and k2, which also introduced a DraIII restriction site at both ends of the 900 bp km cassette. The three fragments were digested using DraIII restriction enzyme (Thermo Fisher Scientific), ligated for 30mn using T4 DNA ligase (Thermo Fisher Scientific), and cloned with the CloneJET PCR Cloning Kit (Thermo Fisher Scientific). The various AES were checked by Sanger sequencing. Each AES was then amplified by PCR on a 3 kb fragment using primer X5 and X6 and purified using QIAquick PCR Purification kit (Qiagen). For gene *yrbE4A*, the occurrence of multiple DraIII and Van91I restriction sites in the targeted sequence led us to modify the cloning strategy. Three PCR fragments were generated using primers I1-I2, I3-I4 and ki1-ki2 and purified using QIAquick PCR Purification kit (Qiagen). Equivalent amounts of each fragment were mixed and a fusion PCR was generated using primers I5 and I6 to generate a 3Kb PCR product used as the AES to disrupt gene *yrbE4A*. Each AES was transformed by electroporation into a recombinant H73Rv strain expressing the recombineering system (van Kessel and Hatfull, 2007) from plasmid pJV53H. Transformants were selected on km-containing plates. Ten colonies were randomly picked and analyzed by PCR using primers X1 and kmR, X4 and kmF and X1 and X7. A clone giving the expected PCR profile (Fig. S3) was selected for further experimentation. The pJVH53 plasmid was cured by sub-culturing and by isolating the selected mutant strains. The loss of pJVH53 plasmid was checked by patching the isolated mutants on hygromycin plate. One hygromycin sensitive clone for each construct was retained for further characterization. For complementation of the *mce1D* mutant, we first transferred the Rv165 cosmid (Brosch et al., 1998), which contains the Mce1 region, into the *E. coli* strain DY380. A zeocin resistance cassette was amplified by PCR using primers Z8 and Z9 and plasmid pMVZ621 (kind gift from Dr. Didier Zerbib, TBI - Toulouse Biotechnology Institute) as a substrate and inserted between the XbaI and Eco147i restriction sites of a mycobacterial integrative plasmid derived from pMV361 (Stover et al., 1993) to give pWM430. A fragment containing the *E. coli* replicon, the mycobacterial integrase, *attP* site and the zeocin cassette was then amplified by PCR using primers M1 and M2 from plasmid pWM430 (3.4kb fragment). The M1 and M2 primers exhibit a 100 nucleotides identity with genes *yrbE1A* and *rv0178* respectively and a 20 nucleotides identity with plasmid pWM430. The 3.4kb amplification fragment was then transformed into DY380:Rv165 after induction of *E. coli* recombineering system as described (Sharan et al., 2009). Transformants were selected on LB Agar containing zeocin (50µg/ml). Recombinant plasmids in which a large *M. tuberculosis* fragment going from gene *yrbE1A* to gene *rv0178* was inserted by recombination into the zeocin containing plasmid were identified by PCR amplification using primers 78 and Z3 or Z1 and F7 and DreamTaq Green polymerase (Thermo Fisher Scientific). A plasmid giving the expected PCR amplification profile was retained for further analysis and named pWM431. This plasmid was transferred by electroporation into the *mce1D* mutant and transformant were selected on zeocin containing plates. The parental and recombinant Mtb strains were made GFP-positive by transfer of a mycobacterial replicative plasmid pWM251, derived from pMIP12 (Le Dantec et al., 2001), which carries the streptomycin resistance cassette from pHP45Ω (Prentki and Krisch, 1984) and the *gfp* under the control of the pBlaF* promotor.

### FA uptake assays in axenic cultures

Alkyne-FA import was quantified by a modification of a previously described uptake assay with radiolabeled FAs (Nazarova et al., 2017). For FA uptake assays, bacteria were grown to an OD_600_ of 0.4-0.8 in 7H9 medium supplemented with 0.5% FA-free BSA, 2g/L dextrose, 0.85g/L NaCl and 0.05% tyloxapol (AD enrichment). Cultures were diluted to OD_600_ of 0.1 in 7H9-AD medium, and incubated with 5µM (unless otherwise stated) of alkyne-FAs pre-conjugated to albumin. For competition assays, natural FAs (5-50µM) and alkyne-FAs (5µM) were simultaneously added to Mtb cultures. After an incubation of 1h (unless otherwise stated) at 37°C, bacterial cultures were collected by centrifugation, washed once in 7H9-AD medium and twice in ice-cold Wash Buffer (0.1% FA free-BSA and 0.1% Triton X-100 in PBS), and fixed in 4% PFA. Imported alkyne-FAs were then stained by a click reaction and analyzed by flow cytometry as described below for bacteria isolated from infected macrophages.

### Macrophage infections

For macrophage infection, mycobacteria were grown from frozen stocks to log phase at 37°C in 7H9-OADC. On the day of infection, bacteria were washed twice and resuspended in phosphate buffer saline, pH 7.4 (PBS). Bacteria were dissociated with the gentleMACS dissociator (Miltenyi Biotech) and remaining clumps were broken by passing through a syringe with a 25G needle and decanting the suspension for 10 min. Bacteria in suspension were quantified by spectrophotometry (OD_600_) and diluted in cell culture medium. Where indicated, macrophages were pretreated with 0.1% DMSO as vehicle control, or with the FADS2 inhibitor SC-26196 (iFADS2) for 2h prior to infection. Confluent macrophage monolayers were infected at a multiplicity of infection (MOI) of 2 bacilli per cell (2:1) for 3h, extracellular bacteria removed by washing with warm DMEM or RPMI-1640 before adding BMDM or THP-1 media. For FADS2 inhibition experiments, the same concentration of iFADS2 was also added after phagocytosis. For enumeration of CFU, 2 volumes of water with 0.3% Triton X-100 were directly added to infected macrophages without supernatant removal. After 10 min at 37°C, lysed cells were serially diluted in water and plated on 7H11 agar supplemented with 0.5% glycerol and OADC. CFU were quantified after incubation at 37°C for 14 – 21 days.

### Mouse studies

All animal experiments were performed in agreement with European and French guidelines (Directive 86/609/CEE and Decree 87–848 of 19 October 1987). The study received the approval by the Institut Pasteur Safety Committee (Protocol 11.245) and the ethical approval by the local ethical committee “Comité d’Ethique en Experimentation Animale N° 89 (CETEA)” (CETEA 200037 / APAFiS #27688). Female, 7-week-old C57BL/6J mice were infected by aerosol route with Mtb H37Rv at a dose of 150-200 CFU per mouse with a homemade nebulizer as described (Sayes et al., 2012). One day prior to aerosol infection and 5 days a week thereafter, mice were given 100mg/kg of FADS2 inhibitor SC-26196 by oral gavage. The lyophilized inhibitor was resuspended in 0.5% m/v methylcellulose – 0.025% Tween 20 as described (Obukowicz et al., 1998). At different time points, mice were sacrificed, and lungs and spleen were homogenized using a MM300 apparatus (Qiagen) and 2.5mm diameter glass beads. Serial dilutions in PBS were plated on 7H11 agar supplemented with 0.5% glycerol, 10% OADC, plus PANTA antibiotic mixture (BD Biosciences) for lung homogenates. CFU were quantified after incubation at 37°C for 14 – 30 days.

### Confocal microscopy

BMDM monolayers cultured on glass coverslips in 24-well plates were infected with GFP-expressing strains of Mtb at a MOI of 2:1 for 3h, rinsed and incubated again in BMDM media. At 24h post-infection, 5µM of alkyne-FAs pre-conjugated to 1% FA-free BSA was added to the cells for 1h. This pulse was followed by a 1h chase in DMEM with 10% complete FBS. Infected macrophages were then washed in PBS, fixed in 4% PFA and permeabilized in 0.5% Triton X-100. Internalized alkyne-FAs were then stained by a click reaction using the Click-iT Plus Alexa Fluor-647 Picolyl Azide kit (Thermo Fisher Scientific) according to the manufacturer’s instructions. Nuclei were counterstained with 1µg/mL of DAPI (Sigma-Aldrich), and coverslips were mounted onto glass slides with Prolong Diamond Antifade Mountant (Thermo Fisher Scientific). Cells were imaged with an LSM 700 inverted confocal microscope and Zen Imaging software (Zeiss), and image analysis was performed using the Icy opensource platform (http://www.icy.bioimageanalysis.org) (de Chaumont et al., 2012). For quantification of FA uptake, all acquisitions were performed using the same settings, mean fluorescence intensity was measured in regions of interest corresponding to BMDMs or bacteria as defined with the HK-Means plugin of the Icy software.

### Flow cytometry quantification of FA uptake

BMDM monolayers were infected with GFP-expressing strains of Mtb at a MOI of 2:1 for 3h, rinsed and incubated again in BMDM media. At 1- or 3-days post-infection, 5µM of alkyne-FAs pre-conjugated to 1% FA-free BSA was added to the cells for 1h. This pulse was followed by a 1h chase in DMEM with 10% complete FBS. BMDMs were harvested, fixed in 4% PFA and permeabilized with 0.5% Triton X-100 in PBS, before staining by click reaction as described above and FACS analysis.

For the analysis of FA uptake by intra-phagosomal bacteria, these were isolated using a previously described procedure (Nazarova et al., 2018). After 2 washes in PBS-0.05% tyloxapol and 1 wash in Wash Buffer, isolated bacteria were fixed in 4% PFA. Fixed bacteria were then permeabilized by incubation with 0.25% Triton X-100 in PBS for 15min at room temperature, and a click reaction was performed as described above. Stained bacteria were washed 3 times in ice-cold Wash Buffer, and resuspended in 7H9 plus 0.85g/L NaCl and 0.05% tyloxapol. Flow cytometry data were collected on a CytoFLEX flow cytometer and analyzed using FlowJo software.

### FA quantification by gas chromatography

BMDMs monolayers cultured in 100mm cell culture dishes (7-9×10^6^ cells per dish) were infected as described above at a MOI of 2:1 with mycobacteria, or treated with 100ng/mL of LPS or Pam3Csk4. At indicated time points, cells were washed with ice-cold PBS (without Mg^2+^ and Ca^2+^), and harvested in ice-cold methanol/5mM EGTA (2:1 v/v). For normalization, cellular DNA content was determined using the Quant-iT dsDNA Assay Kit (Thermo Fisher Scientific) as previously described (Dennis et al., 2010). Respectively 75%/25% of the sample was used for free/total FA extraction using heptadecanoic acid/glyceryl triheptadecanoate (Sigma-Aldrich) as internal standards. Lipids were extracted according to Bligh and Dyer (Bligh and Dyer, 1959) in dichloromethane/methanol/water (2.5:2.5:2, v/v/v), doped with 2µg of suitable internal standard. After vortex and centrifugation at 2000g for 10 min, the organic phase was dried under nitrogen and, for free FAs, directly trans-methylated with 1mL of 14% boron trifluoride (BF3) in methanol and 1mL of heptane for 5 min at room temperature. Total FAs were first hydrolyzed with 0.5M potassium hydroxide (KOH) in methanol for 30min at 50°C, and trans-methylated in BF3-methanol plus heptane for 1h at 80°C. Methylated free FAs were extracted with hexane/water (3:1) as described previously (Stuani et al., 2018), dried and dissolved in ethyl acetate. One microliter of methylated free or total FAs was analyzed by gas chromatography (GC) on a Clarus 600 Perkin Elmer system using FAMEWAX RESTEK fused silica capillary columns (30m x 0.32mm, 0.25µm film thickness). Oven temperature was programmed from 110 to 220°C at a rate of 2°C/min, and the carrier gas was hydrogen (0.5 bar). Injector and detector temperatures were set at 225 and 245°C, respectively. Peak detection and integration analysis were done using Azur software. FAs were quantified by measuring their area under the peak and normalizing to the internal standard. These quantities were then normalized by cellular DNA content.

### FADS2 activity assay

FADS2-mediated bioconversion of deuterated LA-d4 or ALA-d14 was assessed as described previously (Varin et al., 2015). For iFADS2, BMDMs were pre-treated for 2h with indicated doses of iFADS2 or vehicle control before addition of 4µM LA-d4 pre-conjugated to FA-free BSA to the cell culture media. To check FADS2 knock-out, THP-1 monocytes (6.10^5^ cells/mL) were cultured in THP-1 media supplemented with 4µM ALA-d14 pre-conjugated to FA-free BSA. In both cases, after 24h, cells were washed twice in ice-cold PBS. BMDMs were harvested in ice-cold methanol/5mM EGTA (2:1 v/v), while THP-1 were centrifuged and dry pellets were stored at -80°C. Lipids were extracted as described above, total FAs were hydrolyzed with KOH (0.5M in methanol) at 50°C for 30min, and derivatized for 20min at room temperature with 1% pentafluorobenzyl-bromide and 1% diisopropylethylamine in acetonitrile as described (Stuani et al., 2018). Samples were dried, dissolved in ethyl acetate and injected (1µL) on a Thermo Fisher Trace GC system connected to a ThermoFisher TSQ8000 triple quadrupole detector using a HP-5MS capillary column (30m x 0.25mm, 0.25µm film thickness). Oven temperature was programmed as follows: 150°C for 1 min, 8°C/min to 350°C then the temperature is kept constant for 2 min. The carrier gas was helium (0.8 mL/min). The injector, the transfer line and the ion source temperature were set at 250°C, 330°C and 300°C respectively. Ionization was operated in negative ionization mode (methane at 1 mL/min) in selected ion monitoring (SIM) mode and 1 µL of sample was injected in splitless mode. The abundance of each fatty acid and their isotopologue containing 4 or 14 deuterium were obtained by integrating gas chromatographic signals using Trace Finder software.

### Lipid mediator profiling

For lipid inflammatory mediators analysis, BMDMs monolayers cultured in 6-well plates were infected with Mtb at a MOI of 2:1. At indicated time points, cell supernatants were collected, methanol was added at a final concentration of 30% and samples were transferred at -80°C. Samples were thawed, supplemented with 2ng per sample of LxA4-d5, LTB4-d4 and 5-HETE-d8 (Cayman Chemicals), and centrifuged at 5000rpm for 15min at 4°C. Cleared supernatants were submitted to solid-phase extraction using Oasis HLB 96-well clusters, and lipid inflammatory mediators were analyzed by LC/MS/MS as previously described (Le Faouder et al., 2013) on an Agilent LC1290 Infinity ultra-high-performance liquid chromatography system coupled to an Agilent 6460 triple quadrupole mass spectrometer (Agilent Technologies) equipped with electrospray ionization operating in the negative mode. Reverse-phase ultra-high-performance liquid chromatography was performed using a ZORBAX SB-C18 column (inner diameter: 2.1 mm; length: 50 mm; particle size: 1.8 µm; Agilent Technologies) with a gradient elution, and quantifications were obtained using Mass Hunter software.

### Immunoblot analyses

THP-1 monocytes were washed in cold PBS and lysed for 15 min in ice-cold lysis buffer (20 mM Tris, 150 mM NaCl, 1mM EGTA, 1mM MgCl2, 1% n-Dodecyl-(β)-d-maltoside, 50 mM sodium fluoride, and a protease inhibitor cocktail, all purchased from Sigma-Aldrich). Protein concentration was quantified with the NanoDrop-1000 Spectrophotometer (Thermo Fisher Scientific). Cell lysates were resolved on NuPAGE Bis-Tris gels and transferred to nitrocellulose membranes (Thermo Fisher Scientific). Protein detections used the following antibodies: FADS1 (#ab126706, Abcam), FADS2 (#PA576611, Thermo Fisher Scientific), GAPDH (#2118, Cell Signaling Technology). Protein complexes were revealed with the ECL Prime detection reagent (GE Healthcare) and chemiluminescence reading on a Fuji LAS-4000 Luminescent Image Analyzer.

### Gene expression analyses

At the indicated time points, infected macrophages were washed once in cold PBS, lysed with Qiazol lysis reagent and total RNA was purified using miRNeasy kit and RNAse-free DNAse set according to the manufacturer’s instructions (Qiagen). RNA quality was checked using the Agilent RNA 6000 Nano Kit on the Agilent 2100 Bioanalyzer (Agilent Technologies). Gene expression was analyzed using the NanoString’s nCounter digital barcode technology, using a Custom CodeSet encompassing 112 genes including genes involved in metabolism and immune responses, and 5 housekeeping genes (Table S1). Reporter probes, sample, and capture probes were mixed together and hybridized at 65 °C overnight. Following hybridization, samples were transferred and processed in the NanoString nCounter Prep Station. Data was collected by the NanoString nCounter Digital Analyzer, quality checked and normalized to the housekeeping controls using nSolver analysis software according to NanoString analysis guidelines. For quantitative RT-PCR analysis, RNA quantity was assessed using the NanoDrop-1000 spectrophotometer and cDNA was prepared with the High-Capacity cDNA Reverse Transcription Kit (Applied Biosystems). Quantitative PCR was performed using Power SYBR Green PCR Master Mix (Applied Biosystems), on an Applied Biosystems StepOnePlus Real-time PCR system and the following conditions: 2 min at 50 °C, 10 min at 95 °C, followed by 40 cycles of 15 s at 95 °C and 1 min at 60 °C. Primer sequences used in this study were: *Fasn* Fwd, 5’-CCCTTGATGAAGAGGGATCA-3’; *Fasn* Rev, 5’-GAACAAGGCGTTAGGGTTGA-3’; *Fads1* Fwd, 5’-TGGTGCCCTTCATCCTCTGT-3’; *Fads1* Rev, 5’-GGTGCCCAAAGTCATGCTGTA-3’; *Fads2* Fwd, 5’-TCCTGTCCCACATCATCGTCATGG-3’; *Fads2* Rev, 5’-GCTTGGGCCTGAGAGGTAGCGA-3’; *Rpl19* Fwd, 5’-TACTGCCAATGCTCGG-3’; *Rpl19* Rev, 5’-AACACATTCCCTTTGACC-3’. Variations in expression was calculated by the 2^-ΔΔCt^ method with *Rpl19* as endogenous control.

### Statistics

The Graphpad Prism 8 software was used for statistical comparisons and graphical representations, except for the heatmap which was done using the nSolver analysis software. The statistical tests used are detailed in figure legends.

## ACKNOWLEDGEMENTS

This study was supported by Institut Pasteur (T.L., C.D.) and INSERM (U1221, C.D.). We thank the Cytometry and Biomarkers (UTechS CB) Technological Platform for help with nCounter FLEX Nanostring use, and the Image Analysis Hub of Institut Pasteur for confocal microscopy data analysis. We gratefully acknowledge the MetaToul (Toulouse metabolomics and fluxomics facilities, www.metatoul.fr) part of the French National Infrastructure for Metabolomics and Fluxomics MetaboHUB-ANR-11-INBS-0010 for lipid analysis. We also thank Prof. Luke Chamberlain (University of Strathclyde, Glasgow, UK) for sharing his expertise of FA labelling by click-chemistry.

---

**Figure S1:**
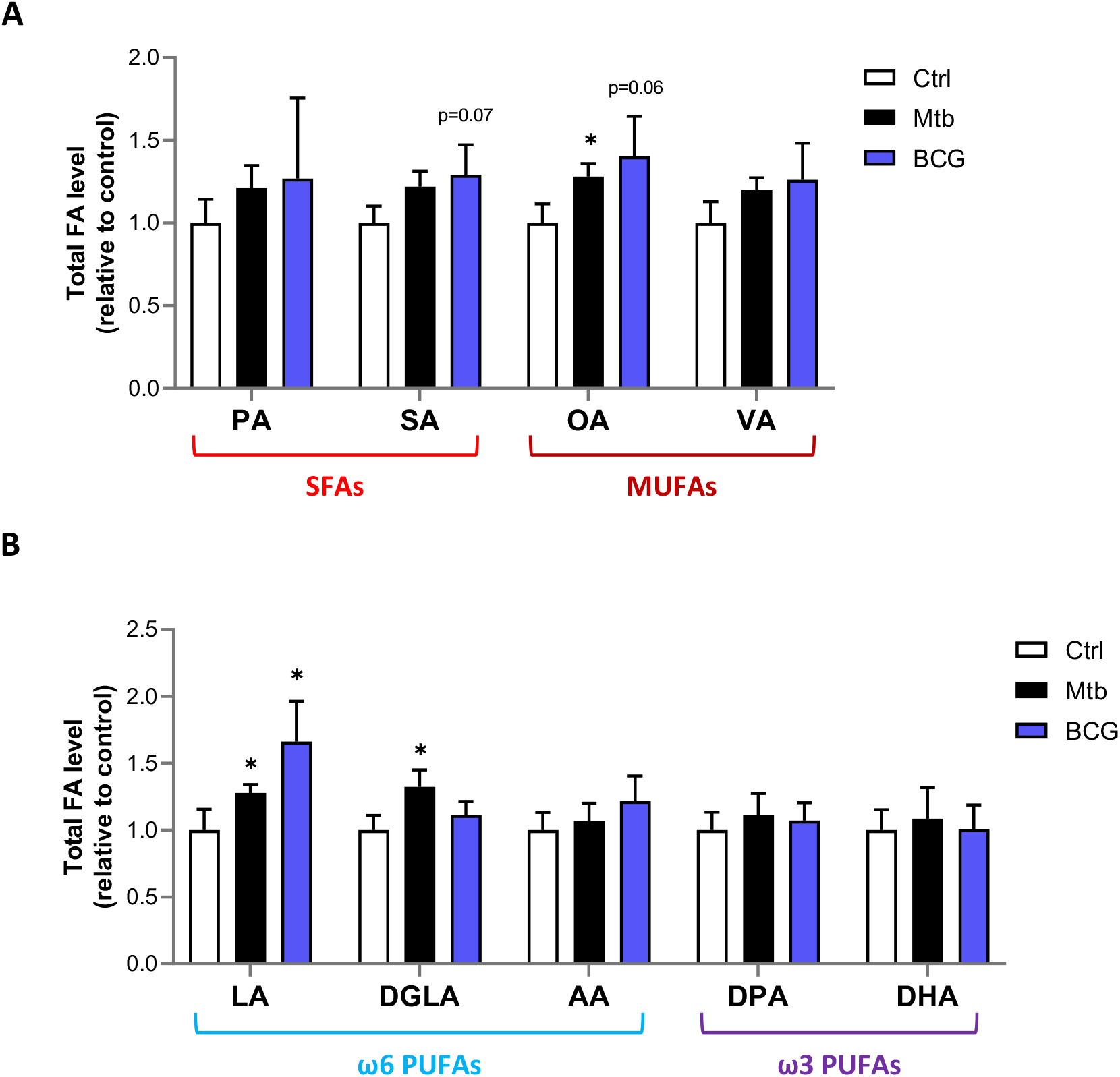
(**A-B**) Intracellular levels of total SFAs and MUFAs (**A**), or total PUFAs (**B**) in BMDMs treated as in (**Fig. 1B**). Data are means +/- SD (n=3) and are representative of 2 independent experiments. \**P*<0.05, unpaired Student’s t-tests.

**Figure S2:**
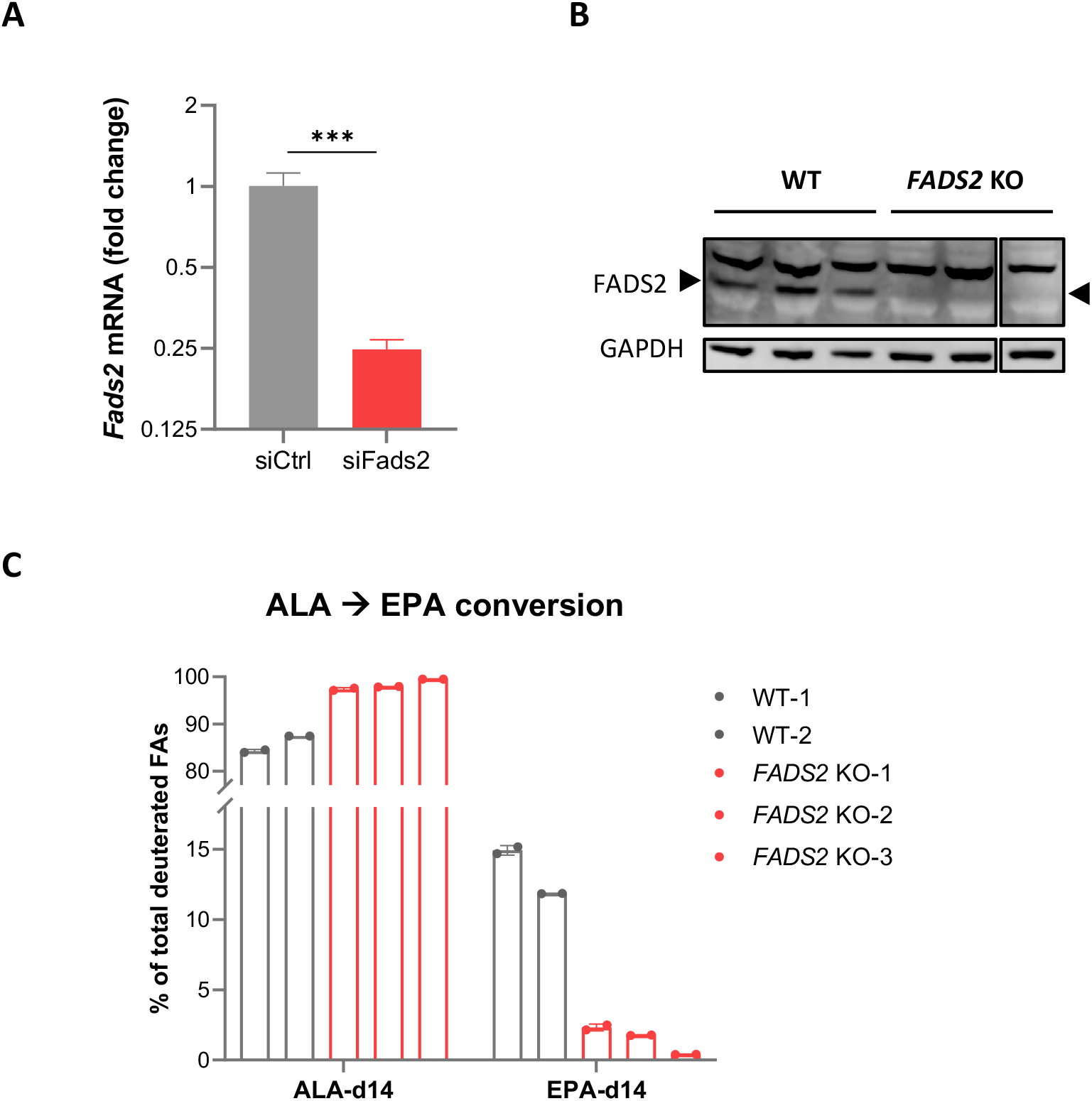
(**A**) Relative mRNA expression of *Fads2* in BMDMs transfected with non-targeting (siCtrl) or *Fads2* siRNA (siFads2) for 48h, as determined by qRT-PCR. Data are means +/- SD (n=3) and are representative of 2 independent experiments. \*\*\**P*<0.001, unpaired Student’s t-test. (**B**) Protein expression of FADS2 and GAPDH (loading control) in wild-type (WT) and *FADS2* knockout (*FADS2* KO) THP-1 clones, as determined by Western blot. Figure S2–Source data 1-3 provides raw blot files (1,2) and a figure with uncropped blot pictures (3). (**C**) Intracellular levels of deuterated ω3 PUFAs in THP-1 clones incubated with the ω3 precursor ALA-d14 for 24h, relative to total deuterated FAs levels. Data are means +/- SD (n=2).

**Figure S3:**
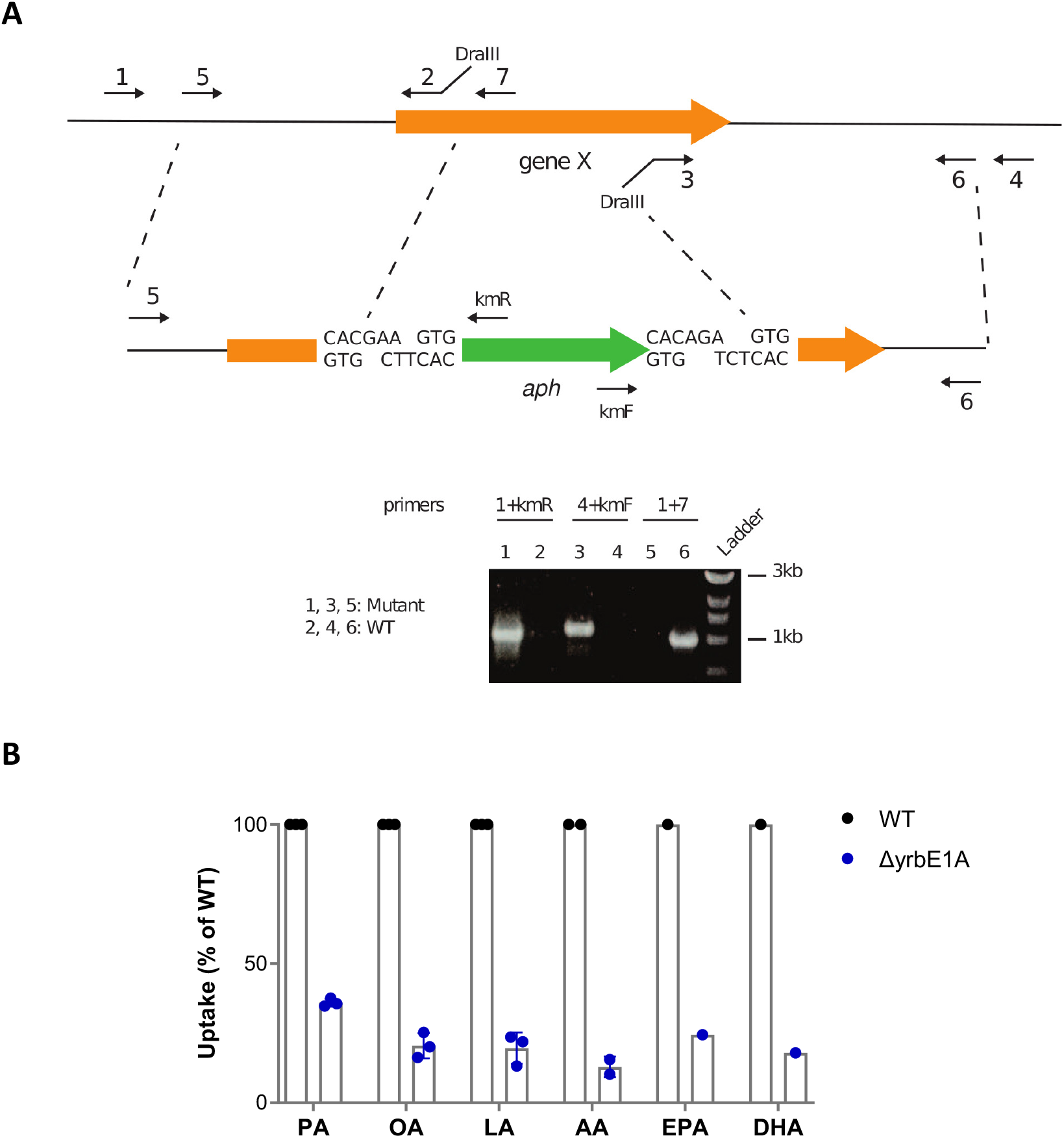
(**A**) Schematics of the strategy used to generate allelic exchange substrates. Figure S3–Source data 1 shows the raw unedited gel. (**B**) Uptake of alkyne-FAs by the ΔyrbE1A mutant of Mtb, relative to Mtb WT. Shown data are means +/- SD (n=1, 2 or 3 independent experiments).

**Figure S4:**
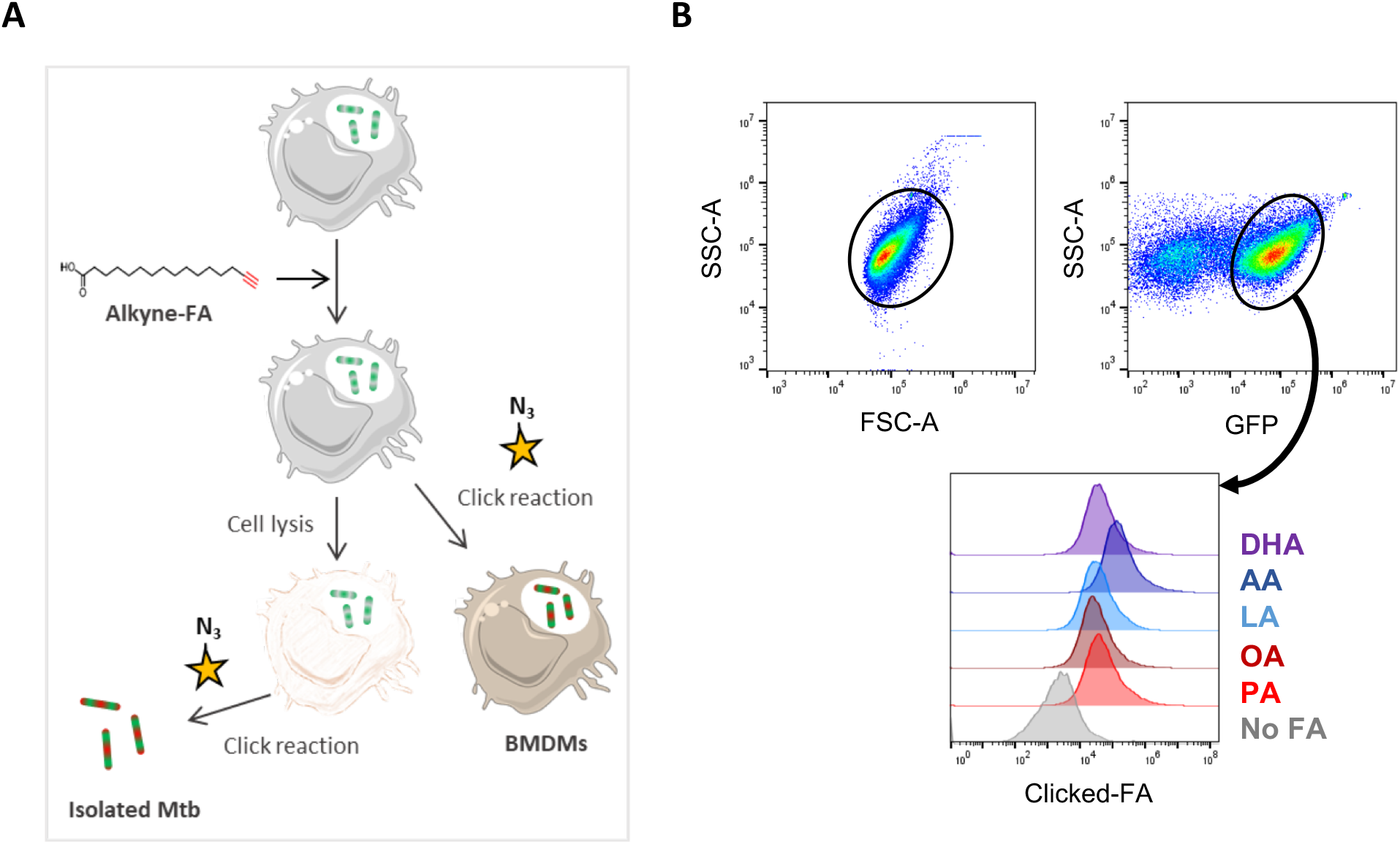
(**A**) Schematics of the click-chemistry approach used to compare the uptake of SFAs, MUFAs and PUFAs by intracellular Mtb and infected BMDMs. (**B**) Gating strategy for the analysis of FA uptake by a GFP-expressing strain of Mtb isolated from infected BMDMs at 24h post-infection, and stained by click-chemistry. Shown data are representative of 3 independent experiments.

**Figure S5:**
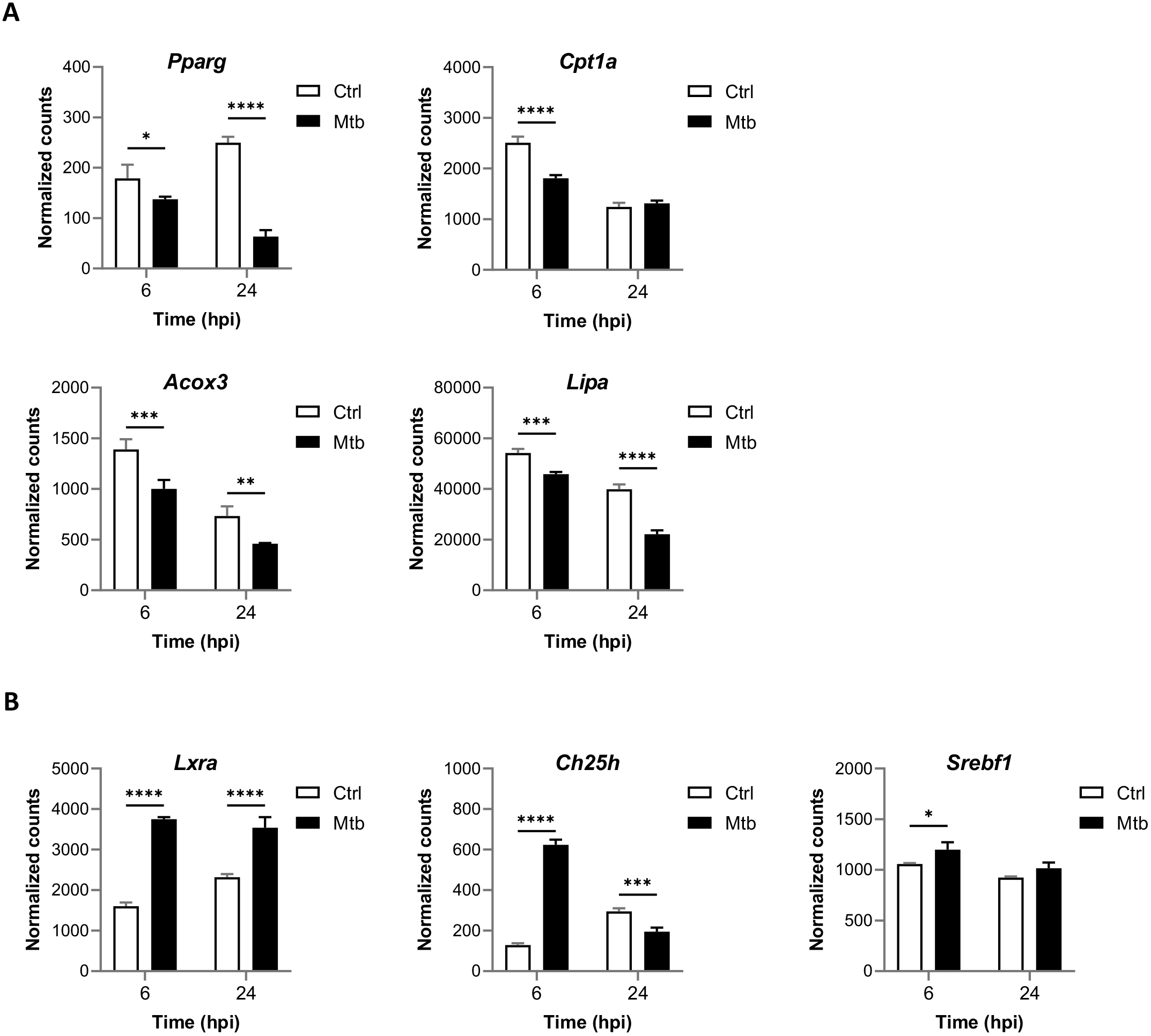
(**A-B**) Relative mRNA expression of PPARγ (**A**) and LXR target genes (**B**), in BMDMs infected with Mtb for the indicated times, as determined by NanoString analysis. Data are means +/- SD (n=3). \**P*<0.05, \*\**P*<0.01, \*\*\**P*<0.001, **** *P*<0.0001 in a two-way ANOVA with Bonferroni post-hoc multiple comparison tests.

**Figure S6:**
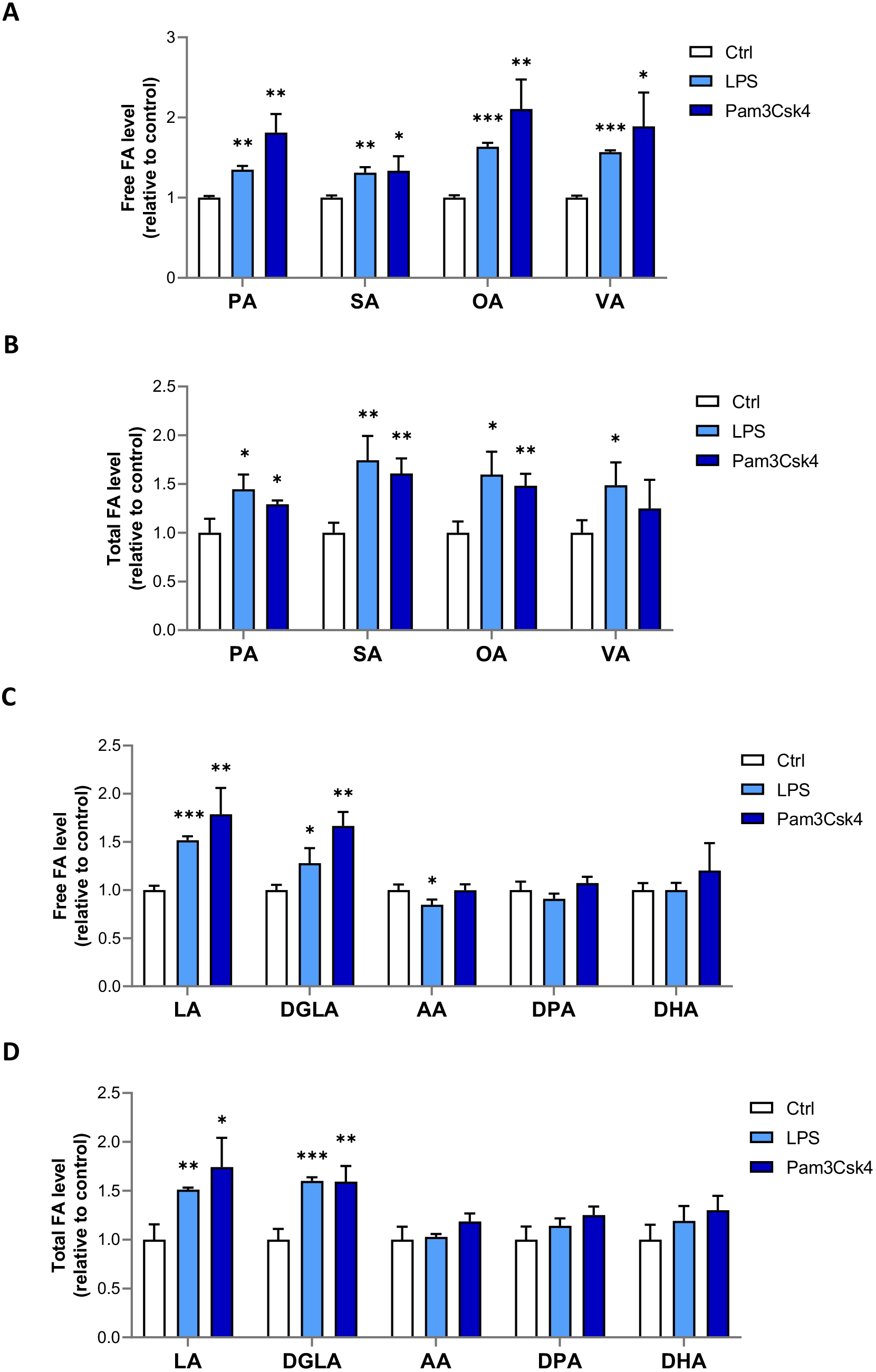
(**A-B**) Intracellular levels of free (**A**) or total (**B**) SFAs and MUFAs in BMDMs treated with LPS (100µg/mL) or Pam3Csk4 (100µg/mL) for 24h. FA levels were normalized to total DNA content, and are shown as fold change relative to Ctrl. (**C-D**) Intracellular levels of free (**C**) or total (**D**) SFAs and MUFAs in BMDMs treated as in (**A**). Data are means +/- SD (n=3) and are representative of 2 independent experiments. \**P*<0.05, \*\**P*<0.01, \*\*\**P*<0.001, unpaired Student’s t-tests.

**Table S1:**
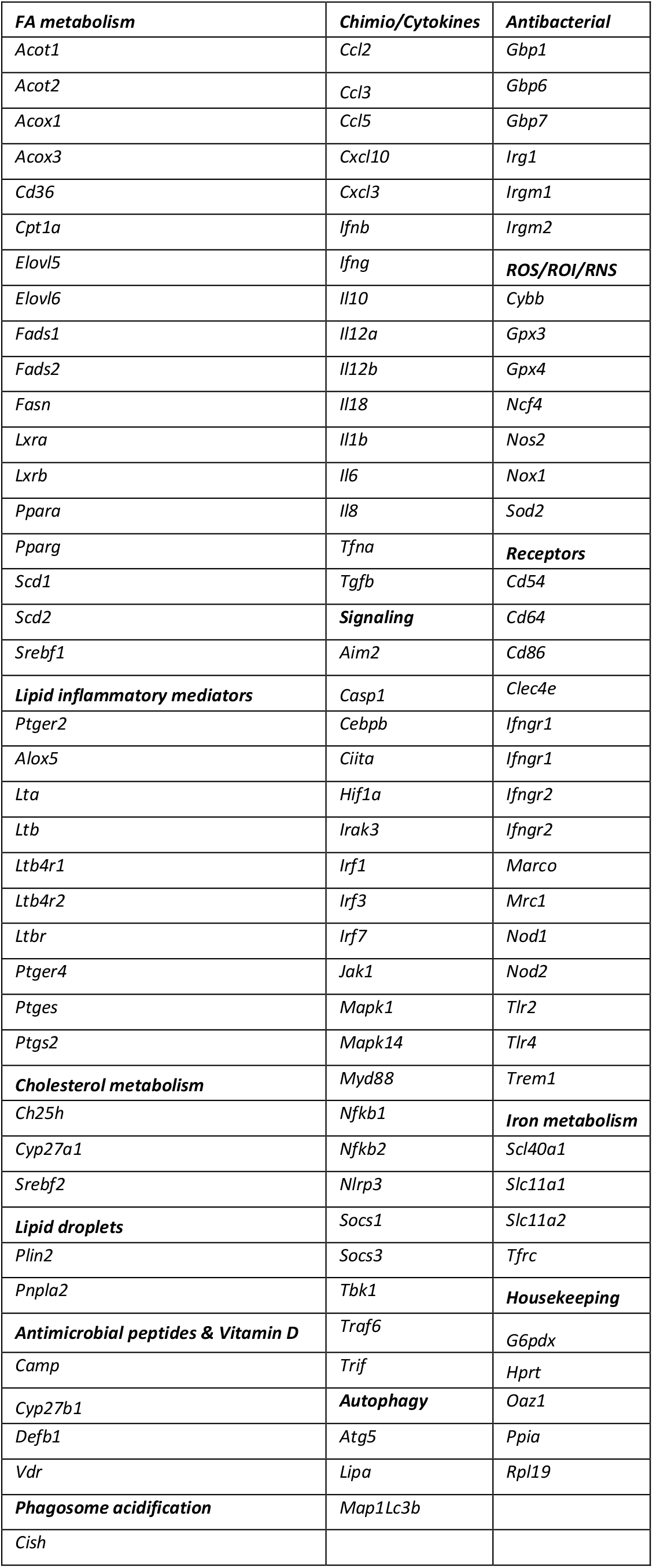
NanoString nCounter Custom CodeSet.

**Table S2:**
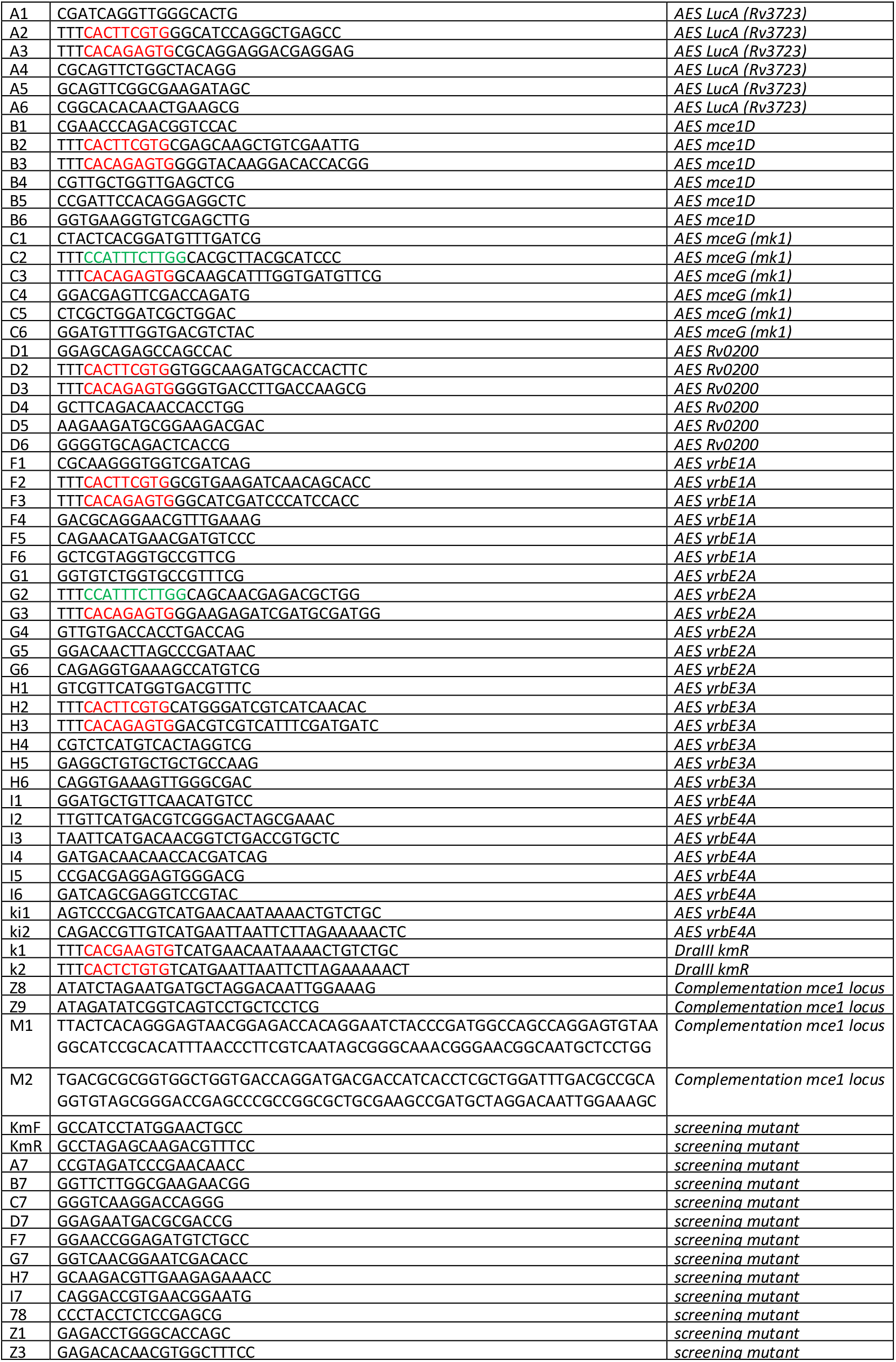
Primers used to generate the various allelic exchange substrates (AES) and Mtb mutants. DraIII and Van91I restriction sites introduced into the primers are indicated in red and green respectively.

